# Structure-guided compound prioritization strategy for virtual screening identifies putative binders for the nuclear receptor LRH-1

**DOI:** 10.64898/2026.06.04.730240

**Authors:** Ana C. Chang-Gonzalez, Alexis N. Campbell, Eric W. Bell, Raymond D. Blind, Jens Meiler

**Affiliations:** Dept. Chemistry, Vanderbilt University, Nashville, Tennessee, USA; Center for Structural Biology, Vanderbilt University, Nashville, Tennessee, USA; Dept. Biochemistry, Vanderbilt University Medical Center Nashville, Tennessee, USA; Dept. Medicine, Division of Diabetes, Endocrinology & Metabolism, Vanderbilt University Medical Center Nashville, Tennessee, USA; Dept. Pharmacology, Institute for Chemical Biology, Center for Applied Artificial Intelligence in Protein Dynamics, Vanderbilt University, Nashville, Tennessee, USA; Institute for Drug Discovery, Institute for Computer Science, Wilhelm Ostwald Institute for Physical and Theoretical Chemistry, University Leipzig, Leipzig, Germany; Center for Scalable Data Analytics and Artificial Intelligence ScaDS.AI, School of Embedded Composite Artificial Intelligence SECAI, Dresden/Leipzig, Germany

**Keywords:** Structure-based drug discovery, computer-aided drug design, virtual screening, hit prioritization, data augmentation, multi-layer perceptron, nuclear receptor, fluorescence polarization

## Abstract

Compound ranking in structure-based virtual screening notoriously yields highly ranked false positive binders due to variable poses or biases in scoring terms. We developed a compound prioritization strategy that utilizes sampled docked poses from contrasting docking approaches (targeted physics-based docking and blind docking with a generative model) against multiple models of the target protein to train a multi-layer perceptron (MLP). The model predicts binders at the orthosteric ligand-binding pocket of the nuclear receptor LRH-1 (NR5A2). Our approach circumvents the reliance on a single docked pose for scoring compounds or individual scoring metrics for compound ranking. In a separate benchmarking set, we observed that the MLP identifies known binders that are chemically dissimilar from the compounds in the training set and is sensitive to single scaffold modifications, making it a potential tool for lead optimization. We applied our strategy to a prospective virtual screening campaign, which resulted in the discovery of four putative LRH-1 binders. We found that a combination of scoring and prediction metrics enriches for the hit compounds across library sizes. In all, this implementation presents a method to leverage structural and experimental data to aid virtual screening for a challenging protein target.

## Introduction

The Liver Receptor Homolog-1 (LRH-1, NR5A2) is a ligand-regulated nuclear receptor expressed in the liver, pancreas, and gut, and a candidate therapeutic target for cancer^1,2^ and metabolic and inflammatory disorders^3–8^. There are over 20 published structures determined experimentally by X-ray crystallography of the ligand-binding domain (LBD) of LRH-1 bound to various phospholipids, synthetic small molecules, and one apo model. In the published structures, phospholipids and synthetic small molecules bind to the ligand-binding pocket, a large (∼1600 A^3^) hydrophobic pocket in the LBD^9–11^. All published structures also contain an LXXLL-motif coregulator peptide bound to the activation function 2 (AF-2) domain. Synthetic compounds drive coregulator peptide binding affinity to LRH-1, likely through allosteric networks that induce conformational changes to the AF-2^12,13^. This mechanism is proposed to modulate LRH-1 target gene expression^13^. Given the clinical relevance of LRH-1, significant effort to use structure-guided approaches has resulted in the development of chemical tool compounds^14^, discovery of LRH-1 antagonists^15^, and detailed structure-activity compound studies^12^. Recent fragment-based screens identified fatty acid mimetic compounds as LRH-1 modulators^16^ and structure-activity relationship (SAR) studies a fragment lead resulted in the discovery of selective LRH-1 agonists with drug-like properties and dose-dependent anti-inflammatory activity^17^. These studies suggest that the expansive LRH-1 pocket may accommodate previously unexplored chemical space, motivating further efforts to identify new compounds considering the current lack of FDA-approved drugs targeting LRH-1.

Structure-based virtual screening (VS) coupled with make-on-demand libraries that boast billions of synthesizable compounds present a cost-effective method to scale screening efforts against the LRH-1 LBD and explore the expansive chemical space^18–22^. LRH-1 structures indicate that phospholipids embed their hydrophobic acyl chains deep in the LBD pocket, while charged headgroups remain solvent-exposed^11,23,24^. Synthetic compounds bound to LRH-1 descend from GSK8470^25^ and share a *cis*-bicyclo[3.3.0]oct-2-ene scaffold which is buried in the LBD pocket^12,26^. Potent descendants contain long (10 carbon) linkers engaging with residues at the mouth of the LBD pocket, mimicking contacts between phospholipids and LRH-1^12,27^. The 6N-10CA compound combines interactions with pocket interior residues forming direct and water-mediated contacts that improve affinity plus hydrogen bonds with residues at the pocket mouth to promote phospholipid-like allostery and coregulator signaling, a pairing that drives LRH-1 gene expression^12^. Separately, computational modeling of LRH-1 binders not based on the GSK8470 scaffold identified from separate screens place compounds either outside of the orthosteric pocket^28^ or in coinciding geometries^17,28^. During preparation of this manuscript, a new structure of the LRH-1 LBD bound to a compound not based on the GSK scaffold was deposited (PDB 9SMQ). The compound makes direct interactions with the LRH-1 pocket interior, bypassing the water-mediated interactions, and is not contacting pocket mouth residues. A structure-based VS campaign against LRH-1 ideally explores interactions across the full spatial context of the binding pocket while accounting for uncertainty in binding location of compounds without crystallographic poses.

Protocols for structure-based VS use computational scores from docked protein-ligand models to rank and prioritize compounds for in vitro or in vivo testing^29–33^. Therefore, compound selection depends on the fidelity of the docked pose. Score functions evaluate the likelihood of compound binding at the target site and are designed to correlate with binding affinity, such that the top-scoring compounds from VS should enrich for hit compounds ^34–36^. However, score functions often include non-binders among top-ranked candidates, resulting in high false positive rates from VS following testing^37,38^. In a direct test case for LRH-1, computational screening using PyRx rigid-body docking was less selective than a fluorescence-based screen to identify novel binders^28^. Efforts to improve score function rankings incorporate machine learning-based adjustments^39–46^, yet these methods often fail to work on proteins or compounds outside of their training sets or in a prospective screening campaign^47,48^.

Here, we present a method to incorporate machine learning with structure-based VS to improve compound prioritization in search of LRH-1 binders. First, we docked compounds from a previous wet-lab screen^28^ to the target pocket using three LRH-1 constructs. Given the large size of the LRH-1 pocket, docking resulted in multiple poses with similarly favorable scores per compound for both binding and non-binding compounds. We extracted individual energy terms accounting for protein-ligand interactions from the docked poses and used the energy terms as features to train two multi-layer perceptron (MLP) predictors. One MLP predicts the likelihood of binding of the compound to LRH-1 and the second which predicts the likelihood of the compound behaving as an activator of LRH-1. By including a variety of poses per compound, training data for the MLP predictors did not solely rely on one docked pose per compound. The docked ensembles helped sample the conformational flexibility in the pocket and possible energetic combinations due to the expansive pocket. Second, we evaluated the trained model on the characterized GSK-based compounds, which are chemically dissimilar to the compounds used for training. We found that the predictions correctly ranked 6N-10CA as the top compound from the set of compounds with structural models determined experimentally by X-ray crystallography, and 6N and 5N as the top compounds from a congeneric series. Finally, we performed a pilot study where we screened 98,000 drug-like compounds and tested 95 compounds prioritized based on the model predictions. In a competitive binding assay, four of the 95 tested compounds were identified as new putative LRH-1 binders (4.2% hit rate). Enrichment of hits within top-ranked compounds was driven by a combination of protein-ligand docking score and machine learning predictions. This approach, made possible by years of structural biology, medicinal chemistry, and wet-lab screening efforts, could be applicable toward accelerating structure-based drug discovery efforts toward other challenging protein targets with substantial experimental data.

## Results

### Multi-layer perceptron trained on docked poses predicts LRH-1 binders

We designed an MLP that uses protein-ligand energy terms from docked poses as input features and outputs the probability that a compound binds to LRH-1 (**Fig. 1**, **Table 1**). The training data was curated from our published screen where we identified 58 compounds from the 2322-compound Discovery Spectrum Library (S2k) that displaced a probe molecule from the LRH-1 orthosteric ligand-binding site^28^. We performed targeted docking using Rosetta^49–52^ of the S2k compounds against LRH-1 to generate docked models of the 58 binders and decoy models of the non-binder compounds. We augmented the true positive entries in the training set by additionally docking only the 58 binders to three LRH-1 structures and using a complementary blind docking protocol DynamicBind^29^ against three LRH-1 structures. Both targeted and blind docking protocols account for protein flexibility and ligand conformer sampling. In all, we generated docked models representing 359 positive (binder) and 1,742 negative (non-binder) labels.

**Figure 1.**
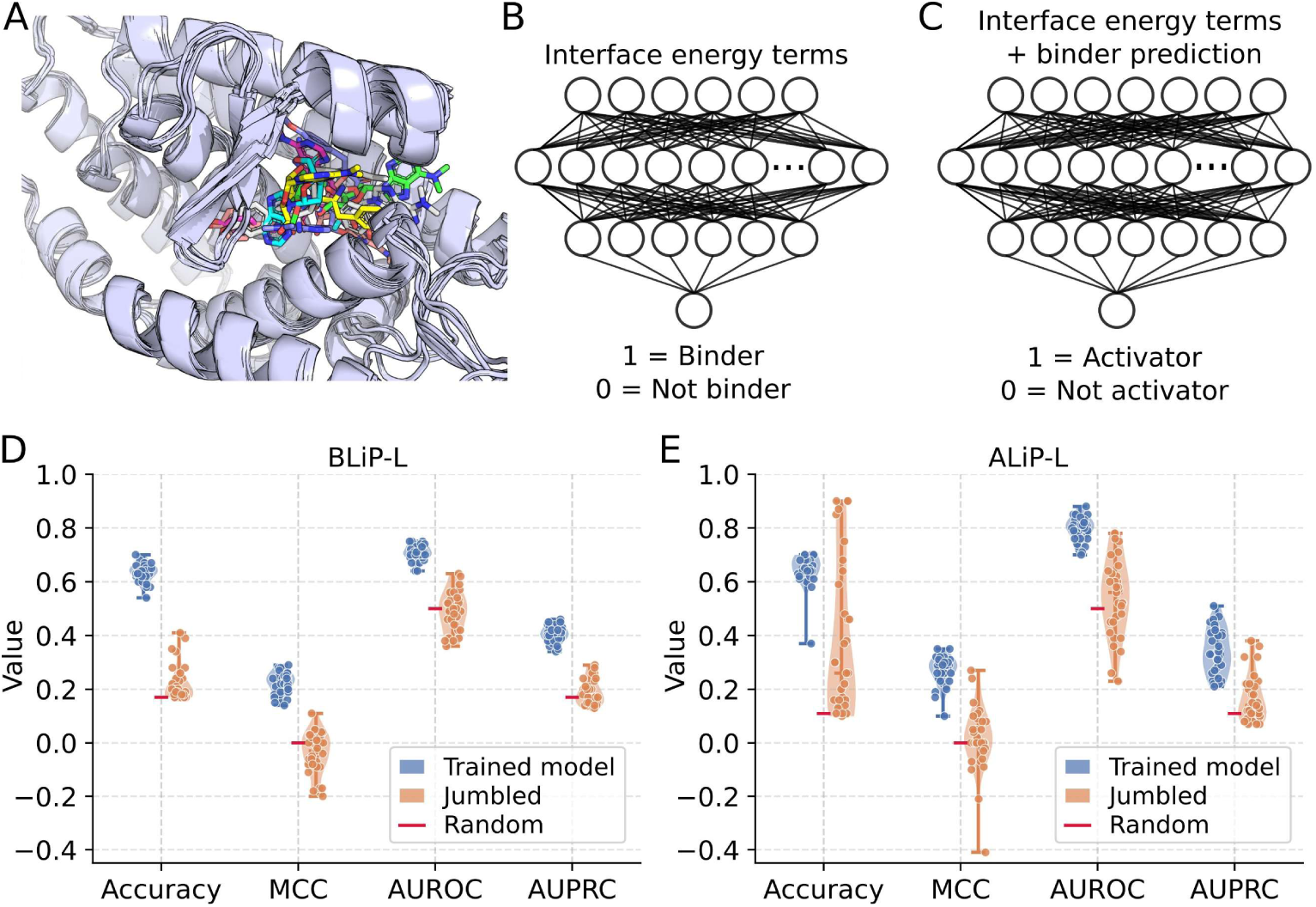
LRH-1 binder and activity likelihood predictors. (**A**) Superimposition of multiple poses from a compound docked to the target LRH-1 (purple) pocket. Seven poses are superimposed from targeted and blind docking protocols. Each pose is rendered in a different color. The poses span the space of the target pocket but differ considerably in orientation. (**B**) Architecture of the MLP used for BLiP-L to predict LRH-1 binder likelihood. The input layer uses six interface energy terms (**Table 1**) derived from the compound poses, as in panel A, of the training set. (**C**) Architecture of the MLP used for ALiP-L to predict LRH-1 activity likelihood. The input layer uses the same six interface energy terms as in panel B plus the binder prediction from BLiP-L. Both models contain two hidden layers and an output layer for binary classification. (**D**, **E**) Performance metrics. Trained model (blue) distributions refer to the trained BLiP-L and ALiP-L models. Jumbled (orange) distributions refer to the baseline model where input features were permuted to disrupt the feature-label relationship. Random (red) line denotes the expected performance for each metric by random guessing. These are: accuracy equal to TP/FP, MCC equal to 0, AUROC equal to 0.5, and AUPRC equal to TP/FP. TP and FP are true positive and false positive, respectively, for the full training set. MCC is Matthews correlation coefficient. AUROC is area under the receiver operating characteristic curve and AUPRC is area under the precision-recall curve. Performance metrics indicate both BLiP-L and ALiP-L distinguish binders and activators, respectively, jumbled features span ALiP-L trained model performance, indicating a less robust feature-label relationship for activity prediction.

**Table 1.**
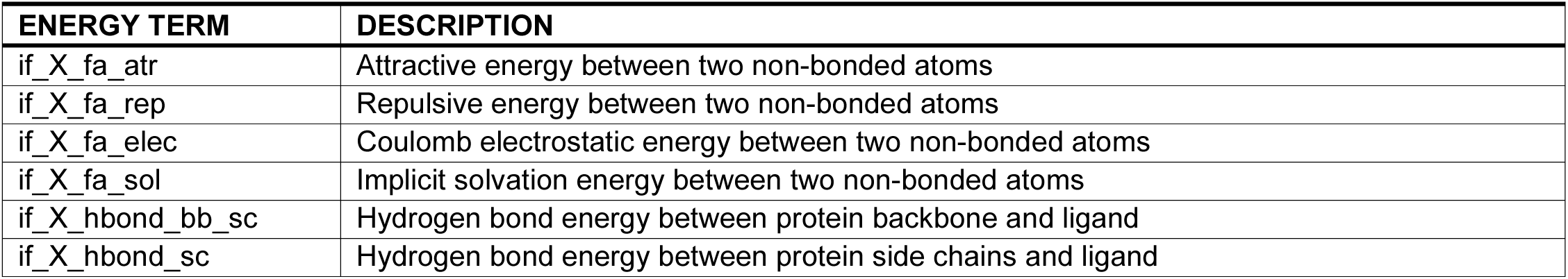
Rosetta protein-ligand interface energy terms used for model input features.

We observed variability in the bound pose of a compound between different docking methods (**Fig. 1A**). Docked compounds engaged residues throughout the target pocket (**Figs. S1A** and **S1B**). Residues most frequently within 3.0 Å of the docked S2k compounds were H390, M345, and L517. H390 in the pocket interior is involved in a π-π stacking interaction with GSK8470 and RJW100-derived compounds^12,26^ or an indirect water-mediated hydrogen bond with compounds 2N, 5N, and 6N^27^, or both as for 6N-10CA^12^, in crystallographic poses. M345 in the deep pocket forms a hydrogen bond with 6N^27^ and 6N-10CA^12^. Mutagenesis and functional studies confirm the relevance of these residues for compound binding^8^. L517 is proximal to the pocket mouth exterior residues involved in interactions with 6N-10CA^12^ and could be involved in nonpolar interactions with LRH-1. In all, the docked models suggest that the S2k compounds interact with LRH-1 residues that are critical for binding. This preference for residue interactions was robust across docking approaches, as we observed similar patterns of high-frequency interaction residues in blind docking poses (**Figs. S1C** and **S1D**).

The cumulative protein-ligand interface score (interface_delta_X or IDX) suggests a difference in the distribution of energy terms between non-binding and binding compounds (**Fig. S2A**). Individual energy terms that compose the cumulative score indicate no single term drives this difference (**Fig. S2B**). The MLP (**Fig. 1B**) enhances the cumulative signal for LRH-1 binder prediction, learning the energetic features that distinguish LRH-1 binders from non-binders. By using energies from multiple docking methods for binder compounds, we account for uncertainty in the docked poses and incorporate the potential for conformational flexibility and different binding modes for new compounds. Cross-validation performance metrics indicate that the MLP can identify LRH-1 binders, capturing feature patterns consistently over a model trained on jumbled features (**Fig. 1D**). The trained and optimized model was saved for future use and named BLiP-L (Binder Likelihood Prediction for LRH-1).

### Separate activity prediction model complements binder prediction

In addition to binder prediction, we developed a separate MLP to predict compound activity to LRH-1. The goal of the model was to distinguish compounds that bind to LRH-1 from ones that both bind and functionally act on the receptor. We curated LRH-1 activity data by using the results from our published S2k screen. In the published screen, we had identified 15 compounds from the 58 binders that either regulated coregulator receptor binding or modulated LRH-1 gene expression in luciferase reporter assays^28^. We used the interaction energy terms from all the docked models as for BLiP-L plus the binder likelihood prediction from the saved BLiP-L model as input features to the activity predictor (**Fig. 1C**). In addition to the data augmentation technique used for BLiP-L, we addressed data imbalance by excluding compounds below the 70^th^ percentile of binder predictions. In all, we used the input features representing 67 positive (activator) and 563 negative (non-activator) labels.

Cross-validation tests demonstrate that the activity prediction model identifies activators with better performance over a random baseline (**Fig. 1E**). However, while BLiP-L cross-validation metrics demonstrate separation between the trained model predictions and that of a model trained on jumbled features, cross-validation for the activity predictor indicates weaker separation, suggesting less robust feature learning. Nonetheless, the trained activity predictor results in a more narrow distribution in performance output, indicating greater consistency in identifying activators compared to a model trained on jumbled features. Therefore, this model is still informative, albeit with lower confidence in its predictions. The trained and optimized model was saved for future use and named ALiP-L (Activity Likelihood Prediction for LRH-1).

### Trained models demonstrate performance on an out-of-distribution dataset

Models trained to predict protein-ligand binding or affinity often fail to generalize to ligands not seen in the training data^39^ in out-of-distribution datasets, where ligands have different chemical or physical properties as the ligands in the training set. Similarity of the S2k compounds to the prototypical RJW100^14,26,27^ compound was quantified by calculating the Tanimoto coefficient of the Morgan fingerprint (**Fig. 2A**). Tanimoto similarity remained below 0.2, indicating the S2k compounds used to train BLiP-L and ALiP-L are chemically dissimilar from RJW100.

**Figure 2.**
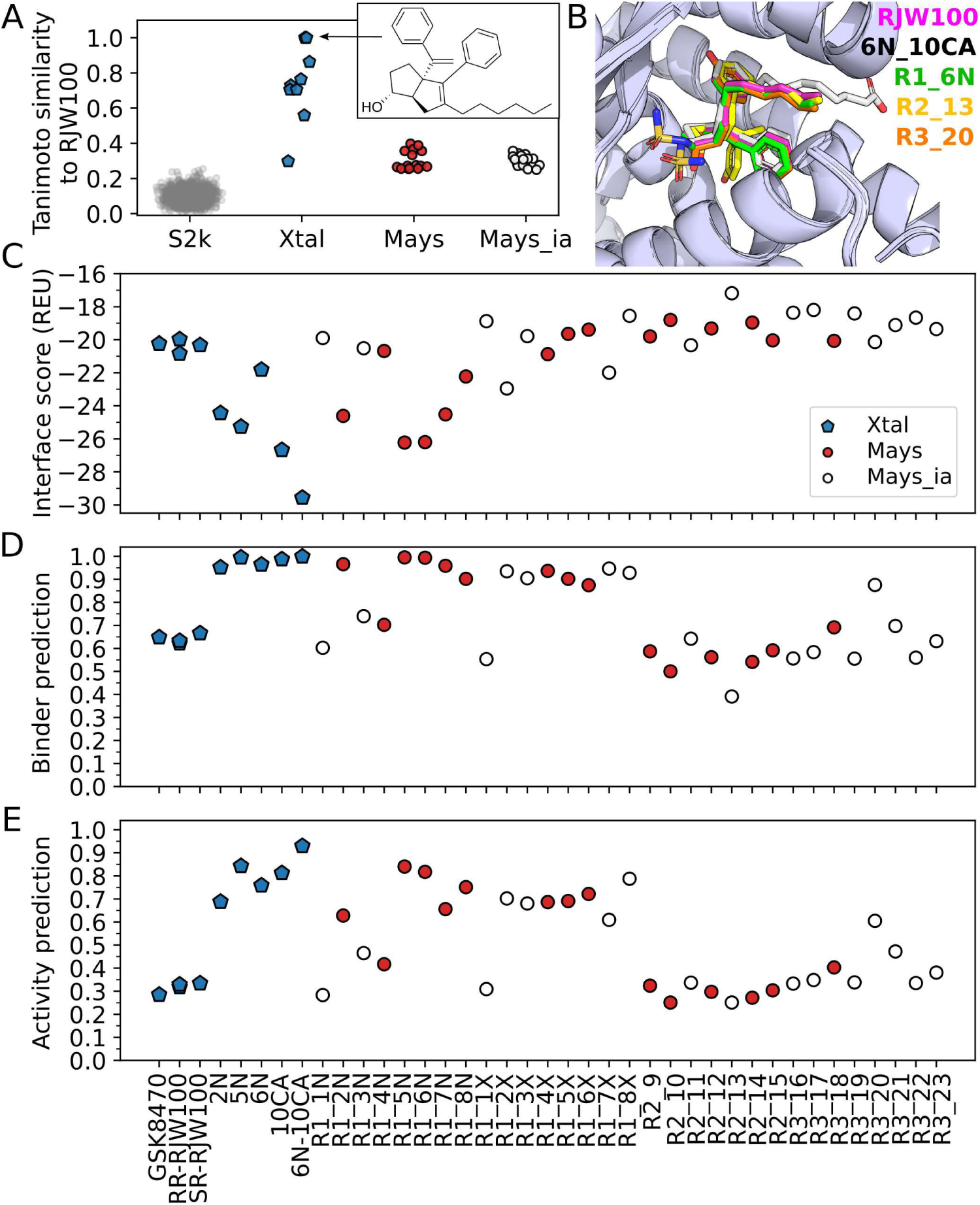
Testing model generalization on out-of-distribution compounds. (**A**) Tanimoto similarity of compounds in different datasets to prototypical RJW100. Values closer to 1.0 indicate higher similarity to RJW100 (inset). Tanimoto similarity was computed with the Morgan fingerprint. S2k are the training set compounds and Xtal are compounds with published structural models (**Table 2**). Mays and Mays_ia are active and inactive compounds, respectively, from a published congeneric series by Mays et al^27^. Xtal and Mays compounds are more chemically similar to RJW100 than S2k compounds. (**B**) Superimposition of crystallographic poses of RJW100 (PDB 5L11), 6N-10CA (PDB 7TT8), and three Mays compounds. Compounds are colored by atom type with standard PyMOL scheme (oxygen red, nitrogen blue, sulfur yellow) and carbon a custom color matching the text color label. The carbon white compound is 6N-10CA. (**C**) Interface score in Rosetta Energy Units (REU) for Xtal and Mays compounds from panel A. (**D**) Binder and (**E**) activity prediction probabilities from BLiP-L and ALiP-L, respectively. Panels **C**-**E** share x-axis labels and legend. Xtal compounds are labeled by compound name with respective PDBs listed in **Table 2**. Mays compounds are labeled by R group replaced and design number corresponding to Figure 2 in^27^. Ranking compounds by score, binder prediction, and activity prediction would all result in the most potent compounds from the respective sets, 6N-10CA from Xtal and R1_6N and R1_5N from Mays, as the top-ranked compounds.

**Table 2.** LRH-1 structural models bound to synthetic compounds used to test prediction models.

| PDB | Bound compound |
| --- | --- |
| 3PLZ | GSK8470 |
| 5L11 | RR-RJW100 |
| 5SYZ | SR-RJW100 |
| 5UNJ | RR-RJW100 |
| 6OR1 | 2N |
| 6OQY | 6N |
| 6OQX | 5N |
| 7JYD | 10CA |
| 7TT8 | 6N-10CA |

There are nine experimentally-determined structures with RJW100 or RJW100-derived compounds in the PDB (**Table 2**). These compounds share a common scaffold and therefore are more chemically similar to LRH-1 than the S2k compounds (**Fig. 2A**, *Xtal*). The compounds bind to LRH-1 in a structurally conserved manner with varied binding affinity^3,12,14,26,27,53^. We calculated the protein-ligand interface score (**Fig. 2C**) and generated the protein-ligand energy terms to apply BLiP-L and ALiP-L to predict binding and activity likelihood of the compounds from the bound poses of the nine structural models (**Figs. 2D** and **2E**). BLiP-L predicted that RJW100 and GSK8470 bind to LRH-1 with binder predictions around 0.6, but ALiP-L predicted low activity likelihood around 0.3. There is an increase in both binder and activity predictions for the next-generation compounds (2N, 5N, 6N, 10CA, and 6N-10CA), where BLiP-L binder prediction was greater than 0.9 and ALiP-L activity prediction greater than 0.7. Both predictions would indicate that the next-generation compounds should be prioritized for experimental testing if nothing was known about them a priori.

Notably, the highest-ranked compound by interface score, BLiP-L, and ALiP-L was 6N-10CA^12^, which is the highest potency synthetic compound from the Xtal list. These results demonstrate that BLiP-L and ALiP-L predictions can be used to identify compounds that are chemically dissimilar to those in the training set and that ranking by predictions could identify potent binders.

### Trained models identify top compounds from a congeneric series

We next evaluated the sensitivity of model predictions to functional group substitutions. For the test dataset, we used the compounds in the published study by Mays et al^27^, where structure-guided rational design was used to create a congeneric series from RJW100 scaffold to explore replacing three R-groups. We refer to this as the “Mays” dataset, which is composed of 23 RJW100-analogs with tested binding and efficacy against LRH-1. Eight analogs replaced the RJW100 R1 hydroxyl, seven replaced the R2 styrene, and eight replaced the R3 phenyl. The full list of compounds is found in Figure 2 of Mays et al^27^. Several modifications did not bind to LRH-1, which we differentiate from active compounds (**“**Mays_ia”). The substitutions render the Mays compounds chemically dissimilar to RJW100 by measured Tanimoto similarity (**Fig. 2A**, *Mays* and *Mays_ia*).

We performed the 23 R-group substitutions in silico, using the bound RJW100 structure as the reference binding mode to generate structural models of the Mays and Mays_ia compounds in the LRH-1 pocket (**Fig. 2B**), calculated interface score (**Fig. 2C**), then applied BLiP-L and ALiP-L to get predictions (**Figs. 2D** and **2E**). The top two highest ranked compounds by interface score, BLiP-L binder, and ALiP-L activity predictions were R1_5N and R1_6N, both of which have sub-micromolar binding affinity to LRH-1 and comparable efficacy to that of RJW100^27^. However, there was no clear cutoff to distinguish the Mays active from Mays_ia inactive compounds. Interface score correlates with reported EC_50_ and efficacy relative to RJW100 by Spearman rank correlation, while predictions suggest only modest non-significant correlations (**Fig. S3**). These results indicate that the three metrics are sensitive to changes in individual chemical scaffolds and can identify promising candidates in a congeneric series. While sorting compounds by interface score would best correlate with functional readouts in one published assay, a combination of the three metrics, interface score and predictions, could be beneficial to prioritize compounds for experimental testing from a virtual screen.

### Compound prioritization strategy results in four putative LRH-1 binders in a prospective screen

We next applied BLiP-L and ALiP-L in a prospective screen. For the pilot study, we performed a virtual screen of the Vanderbilt discovery library, a curated list of 98,765 drug-like compounds (VU98k). We used the BioChemical Library (BCL)^54^ and RosettaLigand^49^ to dock the compounds to three conformations of the target pocket, generated interface energy terms, and used BLiP-L and ALiP-L to predict binding and activity likelihood. We designed sequential filters, first by IDX and prediction cutoffs, then by docked pose location within the pocket, and chemical substructure matching to identify compounds with fatty acid mimetic-like substructures. We excluded compounds that contained PAINS substructures^55^ or failed to pass drug-like filters. We assigned confidence tags (high/medium/low) based on predictions for the reduced list of 11k compounds, then curated a list of 95 compounds that are chemically diverse (**Fig. S4A**) with docked poses that suggest the compounds engage known ligand-binding residues spanning the large LRH-1 target pocket (**Fig. S4B**).

The selected compounds were tested for competitive binding to the LRH-1 target pocket (**Fig. 3A**). We screened 10 μM compound against 100 nM purified LRH-1 isolated LBD with 50 nM rhodamine-PE (Rh-PE) probe using a fluorescence polarization (FP) assay. We used RJW100 as a positive control and a well with only LRH-1 as a negative control. A decline in polarization of the fluorescent signal would indicate that the screened compound displaces the Rh-PE probe and therefore binds to the LRH-1 target pocket. We identified seven out of the 95 compounds that decreased FP signal below baseline with log_2_ negative control minus log_2_ compound wells ≥ 0.5 (**Fig. 3B**). We then tested the seven hits for intrinsic fluorescence at the assay spectra. Three of the seven compounds exhibited fluorescence above control wells at the same wavelength used in the polarization assay, therefore we discarded those compounds as false hits. This resulted in four putative LRH-1 binders discovered from our virtual screen (**Figs. 3C-F**).

**Figure 3.**
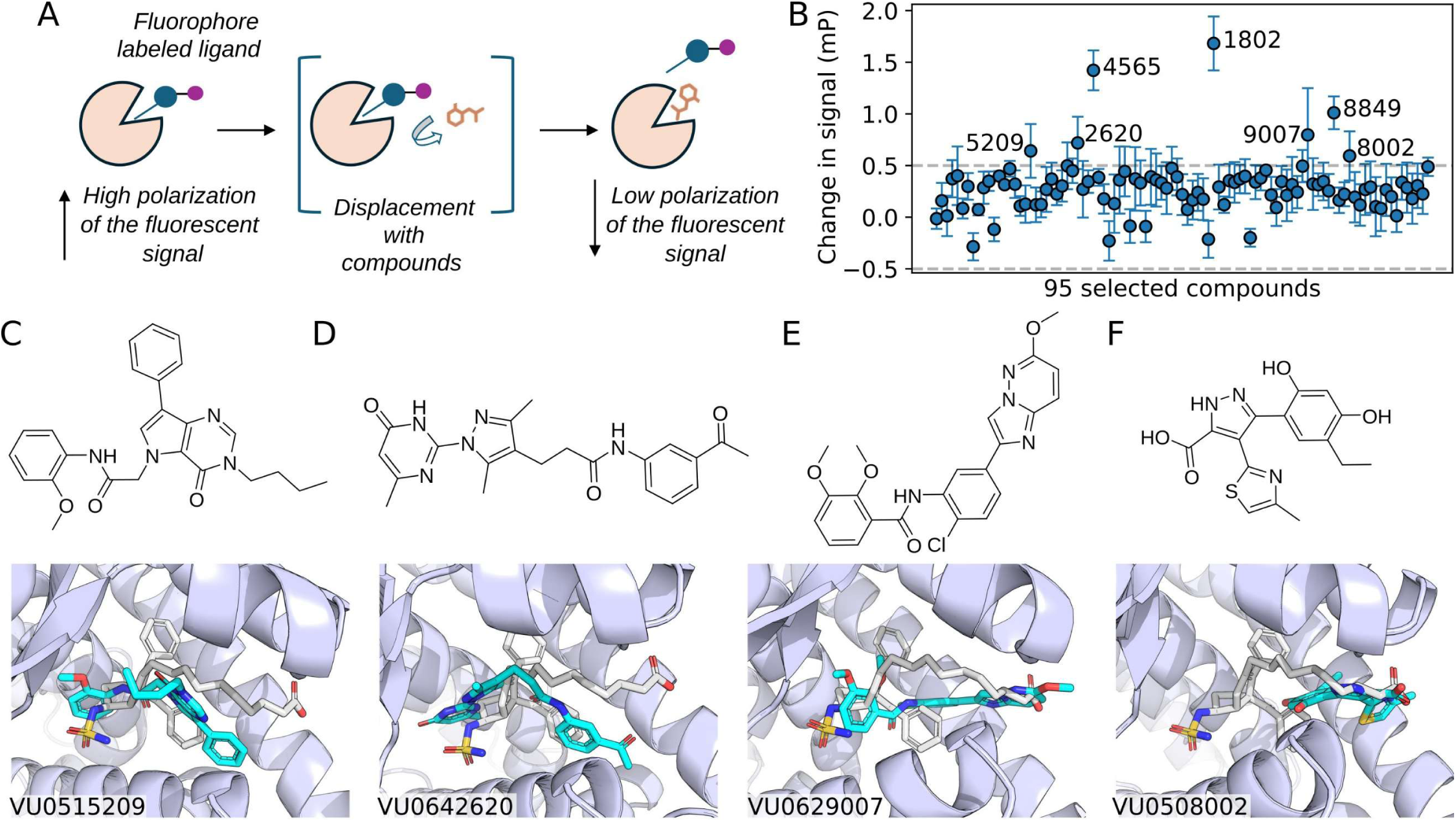
Testing 95 selected compounds from a virtual screen identifies four putative binders. (**A**) Competitive binding fluorescence polarization assay used to test selected compounds. Fluorescence signal changes when a compound displaces the Rhodamine-PE (Rh-PE) probe in the target LRH-1 pocket. (**B**) Log_2_ change in signal computed by log_2_ negative control minus log_2_ compound wells in millipolarization (mP) values induced by 10 μM each of the 95 selected compounds after virtual screen of the VU98k library. Three technical replicates were performed. Seven compounds changed fluorescence intensity by ANOVA multiple comparisons test (p-value < 0.05), labeled in the plot by the last four digits of the VU number identifier. (**C-F**) Four of the seven compounds that passed the intrinsic fluorescence interference screen to exclude potential false positive compounds. Top panel depicts the chemical structure, bottom panel depicts the docked pose of the compound (cyan) with VU number identifier used for scoring and prediction metrics. Docked model is superimposed with crystallographic pose of compound 6N-10CA (white, PDB 7TT8) for reference.

Classically, nuclear receptors are regulated by ligand-induced allostery^56,57^, which alters the interaction between the nuclear receptor and an LXXLL-containing transcriptional coregulator^58–61^. We tested whether the compounds altered coregulator binding using a secondary fluorescence polarization assay with isolated LRH-1 LBD at 10 μM compound, 0.38 μM purified LRH-1 LBD, and 50 nM FAM-labeled DAX-1 peptide. None of the 95 compounds altered LRH-1 LBD binding to either coregulator. In our S2k screen, we discovered compounds that competitively bound to the LRH-1 pocket and regulated full-length LRH-1 gene expression in reporter assays using mammalian cells, but they failed to modify coregulator binding^28^. We proposed a direct inhibition mechanism where the compounds regulate LRH-1 activity by preventing the signaling phospholipids or other molecules from binding to the orthosteric pocket. Our results suggest the four putative binders we identified may regulate LRH-1 in a similar manner. Further functional and structural studies are needed to measure binding and confirm the activity of the discovered compounds.

### Interface score and predictions together enrich top-ranked compounds for LRH-1 putative binders

All four hits resulted in interface scores less than -14 REU, and the minimum binder and activity prediction likelihoods were 0.85 and 0.62, respectively (**Figs. 4A** and **4B**). Visualization of the prediction-score landscape for VU98k compounds indicates that hits were not confined to the expected low-score, high-prediction region (**Figs. 4A** and **4B**). Instead, the hit compounds spanned the range of the landscape covered by the 95 tested compounds. No individual energy term, score, nor prediction metric exhibited clear class separation between tested hits and non-hits (**Fig. S5**).

**Figure 4.**
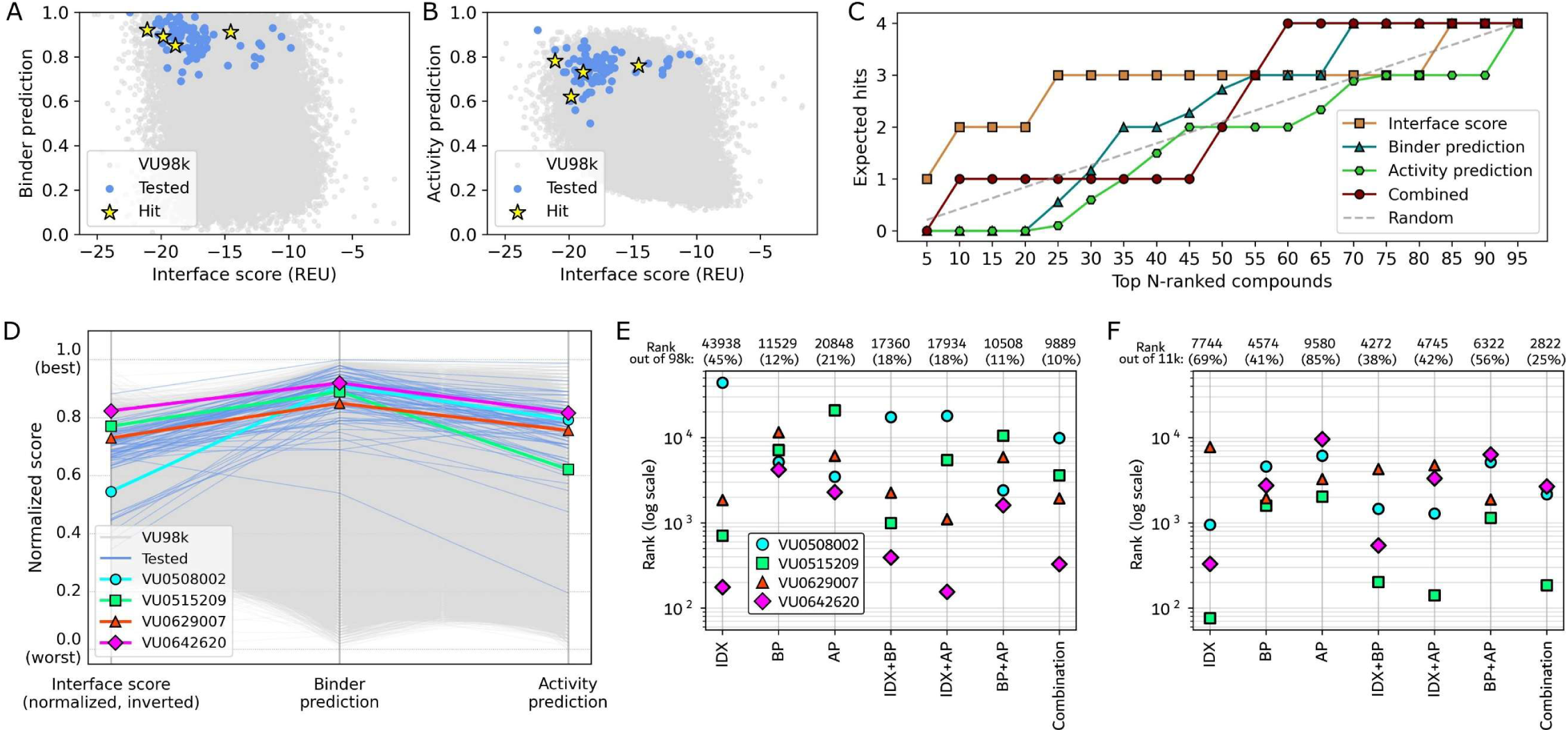
Hit recovery is improved by combining score and prediction metrics across library size. (**A**) Binder prediction versus protein-ligand interface score. (**B**) Activity prediction versus protein-ligand interface score. Gray cloud in panels A and B denote the landscape covered by the VU98k library compounds. Blue cloud denotes landscape covered by the 95 tested compounds from the VU98k set, and yellow stars denote the four hits (**Fig. 3C-F**). (**C**) Number of hits within a set of N-ranked compounds. The 95 tested compounds were ranked by the metric specified in the legend. “Combined” uses the sum of the rankings from the interface score and both predictions as the ranking metric. Tied compound ranks were assigned the average rank. The random baseline denotes the expected hits if compounds were picked randomly without any metric guidance. Expected hits are equal to 4*(N/95). (**D**) Parallel coordinate plot. Each line represents one compound, tracked individually across the three ranking metrics. Gray lines represent all VU98k compounds, blue lines the 95 tested compounds, and colored lines the four hits. Interface score was normalized and inverted such that the lower energy values map to higher normalized scores. Prediction probabilities were used such that the value is normalized to [0,1], where 1 indicates the best predicted value. Ranks of the four hit compounds out of the (**E**) VU98k library and (**F**) reduced 11k compound list across individual and combined metrics interface score (IDX), binder prediction (BP), and/or activity prediction (AP). “Combination” uses all three metrics as in panel C. Rankings for combined metrics use the sum of the individual metric rankings for sorting compounds. Annotated at the top of the plot are the rank and percentage of the library required to capture all four hits per ranking scheme. Lower ranks indicate fewer compounds required for testing to recover all confirmed hits.

To quantify enrichment for hits within top N-ranked compounds per metric, we sorted the compounds by either interface score, binder prediction, or activity prediction, then assigned ranks based on the sorted list (**Fig. 4C**). Ties were assigned the average rank for the set of tied compounds. We also sorted compounds by a combined metric using the sum of the compound ranks from the three metrics. Ranking based on interface score alone recovered three out of the four hits within the top 25 compounds, outperforming the predictions and combined metric in hit enrichment (**Fig. 4C**, *orange* vs others). However, ranking based on the combined metric recovered all four hits within the top 60 compounds (**Fig. 4C**, *maroon*). The combined metric improves prioritization over scores or predictions alone by identifying all four hit compounds within the smallest set of top-ranked candidates.

Expected hit recovery using activity prediction was consistently below the random baseline (**Fig. 4C**, *green*), indicating that ranking compounds by activity prediction is less effective at recovering hits than expected if compounds were randomly ranked. In fact, when the activity prediction is excluded from the combined sum, the combined sum ranking remains unchanged. Parallel coordinate analysis revealed a consistent profile among the four hit compounds where binder prediction increases relative to interface score (**Fig. 4D**). The largest increase was for the hit with the lowest interface score, VU0508002 (**Fig. 4D**, *cyan*). Activity prediction on the other hand introduced a downward deflection across the hit compounds. However, by visual inspection a few of the tested non-hits (**Fig. 4D**, *blue*) that ranked highly by interface score alone demonstrated pronounced decreases in activity prediction. This suggests that the inclusion of activity prediction in the combined rank may help deprioritize compounds ranked highly by interface score alone that are not hits in an experimental assay.

The contribution from predictions in improving hit ranks is apparent when ranking the compounds within the context of the larger 98k and 11k sets. Combination-based ranking results in the lowest maximum rank of all four hits compared to ranking by any of the three individual metrics alone or two-metric combinations for both the 98k (**Fig. 4E**) and 11k (**Fig. 4F**) compound sets. The combination ranking renders all hit compounds within the top 10% of the 98k compounds (**Fig. 4E**) and within 25% of the 11k compounds (**Fig. 4F**). The second best ranking metrics in both sets include at least one of the predictions. These results indicate that integrating interface score and prediction metrics consistently outperforms individual or two-metric combinations for hit prioritization, regardless of library size.

## Discussion

In this work, we leveraged the results from our previously published fluorescence-based screen^28^ to design binder and activity prediction models, then applied the models to guide compound selection in a prospective virtual screen. Utilizing protein-ligand binding scores and predictions, we prioritized and selected 95 out 98k compounds to test in a competitive binding assay. Four out of the 95 tested compounds were identified as new putative binders for LRH-1, yielding a 4.2% hit rate (4/95 tested compounds). We found that combining the docked pose protein-ligand interface score with both binder and activity predictions enriches the top-ranked compound list for the four putative binders.

Ranking compounds from a virtual screen by in silico metrics alone commonly results in highly ranked false positives^62–70^. Therefore, after initial threshold-based filtering, our strategy for prioritizing compounds for testing avoided selecting only the top-ranked compounds by a single ranking metric. While we cannot state whether the non-tested compounds from the VU98k library would bind to LRH-1, our selection strategy allowed us to test 95 compounds that covered a wide range of values for scoring and prediction metrics, sampling the metric landscape and therefore providing data to explore how individual and combinations of metrics improve ranking hit compounds. Notably, we found that top-ranking compounds by one metric were not consistently prioritized by other metrics, highlighting the potential value of integrating complementary information with multiple metrics for hit prioritization (**Fig. 4**). Toward this, we found that ranking compounds in expanded libraries using combinations of metrics lowered the maximum rank of hit compounds, indicating that in a hypothetical screen, fewer compounds would need to be tested in the wet-lab to identify hits (**Figs. 4E** and **4F**).

Chemical and structural filtering were used in addition to rank-based filtering to prioritize compounds for testing. Retrospective analysis of the tested compounds suggested limited discriminative value of the chemical substructure filtering for compounds with fatty acid mimetic-like substructures (**Table S1**). While individual motifs appeared in the hit compounds, these features were also present among non-hit compounds, and no single checked motif was consistently enriched across all hits. Although hits tended toward lower minimum distances to selected LRH-1 pocket residues, they were not clearly separated from the distribution of tested non-hits (**Fig. S6A**). This indicates that the structural filter was useful for restricting the search space to a set of compounds expected to dock proximally to the target pocket, but provides limited additional discriminative power within a pre-filtered subset. Both chemical and structural filters were restricted to a predefined set of criteria. We limited chemical substructure filtering to a predetermined set of motifs and did not systematically explore alternative motif definitions nor related chemotypes. Likewise, we limited structural filtering to a predetermined set of pocket residues. Such guided filtering may be useful when chemical motifs or interacting residues are known for a target a priori, but would reduce chemical diversity when motif definitions are incomplete or when residues that determine binding or activity extend beyond spatial proximity.

Docking against the three receptor conformations was used to incorporate variability in the local LRH-1 pocket environment and improve sampling of plausible binding modes. The most favorable docked pose for the four hit compounds was not consistently associated with a single construct. Rather, two hits favored the model based on the small-molecule bound pocket and the other two hits favored the phospholipid-bound pocket (**Fig. S6B**). While the docked poses varied across constructs, the four hit compounds satisfied the applied distance cutoff across the three constructs, suggesting compatibility with the targeted binding region independent of any single docking model. In contrast, 14 of the 95 tested compounds had at least one construct-specific pose that failed the distance cutoff criteria, suggesting that persistence of the structure filtering criteria across receptor conformations could provide an additional metric for compound prioritization. In fact, the success of RosettaVS is partially attributed to the ability to model receptor flexibility in their virtual screening protocol^33^.

Our approach is most useful in identifying likely binders from a large library of compounds over determining the precise affinity order of compounds, as we observed by evaluating score and prediction metrics on the RJW100-derived benchmark set of compounds and congeneric series (**Figs. 2** and **S3**). The score and ranking metrics are able to consistently identify the experimentally-favored compounds from the congeneric series (**Figs. 2C-E**) even though the docked poses are nearly identical (**Fig. 2B**). This sensitivity suggests that the metrics could be used in a hit-to-lead optimization step. Once a weak binder is found, an in silico construction of a panel of modifications as suggested by medicinal chemists that are synthetically accessible and would yield appropriate pharmacokinetic properties could be designed, then evaluated by the scoring and prediction metrics to identify and prioritize leads for testing.

Confidence and compound selection thresholds were empirically determined using benchmarking predictions of the RJW100-based compounds. Screening, docking, and scoring metrics perform differently across protein targets and assays of interest^71^. Machine learning-based predictions often don’t generalize to new targets or chemically different compounds beyond those used for model training^72,73^. The large, flexible, and hydrophobic LRH-1 pocket presents a particularly challenging target for structure-based drug discovery, as compounds can dock in a variety of modes that complicate compound scoring and ranking, and therefore selection for testing in the wet-lab. Consistent with broader limitations in predictive model transferability, generalized scoring functions and machine learning approaches likewise may underperform on targets with chemically and structurally complex pockets. Our approach leverages pose diversity sampled across docking strategies and multiple states in a target-specific framework, enabling the discovery of new putative binders for LRH-1 (**Fig. 3**) and indicating scalability across library size (**Fig. 4**).

Although this work focused on LRH-1, our approach can be applied to other proteins. The prediction models can be retrained using binding and activity data for a different target protein, and the combined scoring and prediction metrics can be used to prioritize compounds for testing and improve hit enrichment. However, care is necessary to ensure experimentally-backed data for training. Similarly, a major limitation of this work is in the fact that we were unable to identify compounds that modified coregulator binding to LRH-1. While these compounds may still regulate LRH-1 target gene expression^28^, since the activity prediction model was trained using compounds that would modify coregulator binding, we expected that some of our hits should modify coregulator binding. This could be due to insufficient true positive samples for training, as only five of the 58 compounds in our training set modified coregulator binding in LRH-1^28^. Also highly likely is that this is due to a limitation in the features used training. In a biological system, compound binding to a target is not always directly related to function. Other factors, such as allosteric effects, receptor dynamics, cofactor recruitment, or external biological implications may contribute to activation. Future work could involve feature engineering to incorporate these effects.

This work demonstrates integration of computational methods to inform experimental testing. Our approach is robust to chemically dissimilar compounds, allowing for wide chemical space searches, and maintains hit enrichment across library sizes. This could be powerful in the context of ultra-large library screening, where prioritizing compounds amid high false positive rates when screening billion-scale libraries is critical^19,22^. Strong retrospective benchmark performance and success finding hits in a prospective screen highlight the potential of this approach to help accelerate drug discovery efforts.

## Figures and Tables (Supplementary Material)

**Figure S1.**
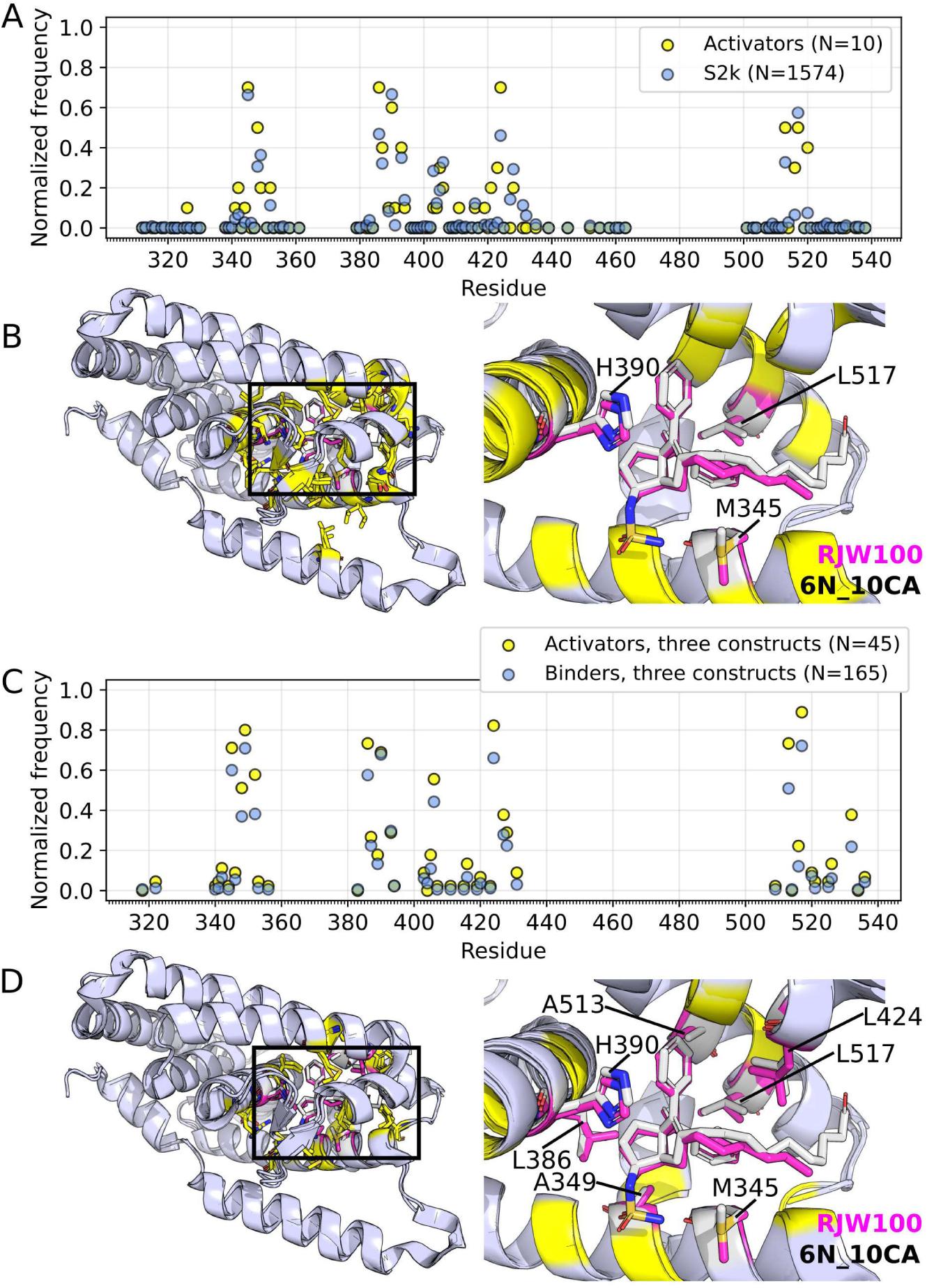
Docked S2k compounds interact with known LRH-1 ligand-binding residues. Normalized frequency of ligand-LRH-1 residue contacts identified using CPPTRAJ native contact analysis from (**A**) targeted and (**C**) blind docking structures. Contacts were defined as non-hydrogen atom interactions within 3.0 Å between the compound and select LRH-1 residues. Poses without contacts were excluded. Number of structures per label indicated in the legend. Docked poses from docking to three LRH-1 models were included in panel C. Residues with greater than 0.5 interaction frequency with compounds from (**B**) targeted and (**C**) blind docking structural models mapped onto models of RJW100 (PDB 5L11) and 6N-10CA (PDB 7TT8) in the target pocket.

**Figure S2.**
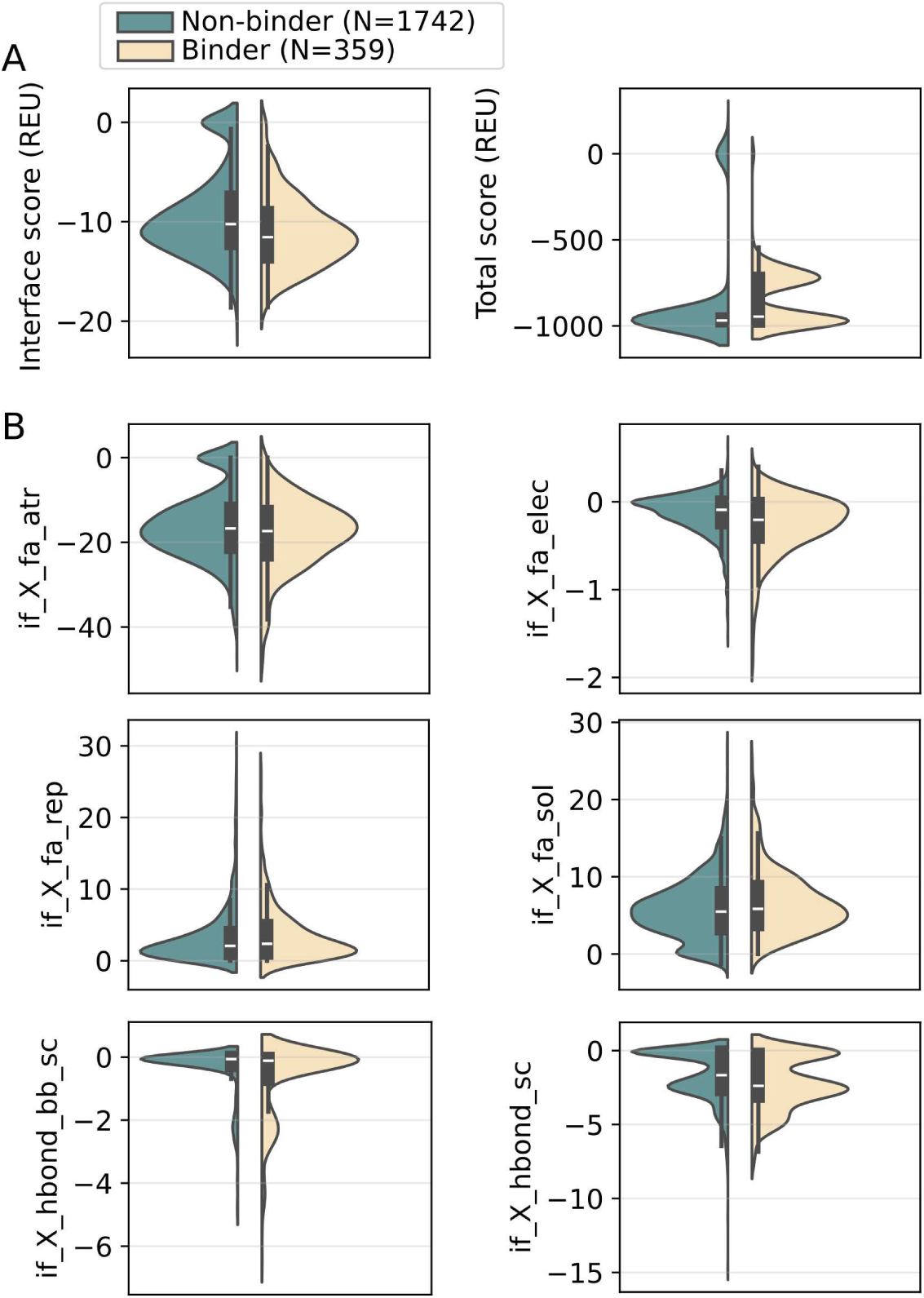
**Docked pose protein-ligand terms compared to compound class**. Violin plots represent the distribution of non-binder versus binder compounds for RosettaLigand (**A**) cumulative scores and (**B**) individual energy terms from docked poses of the S2k library compounds. Box plots indicate the interquartile range, dataset median (white line), and range. The number *N* of compounds included in each class is annotated in the figure legend.

**Figure S3.**
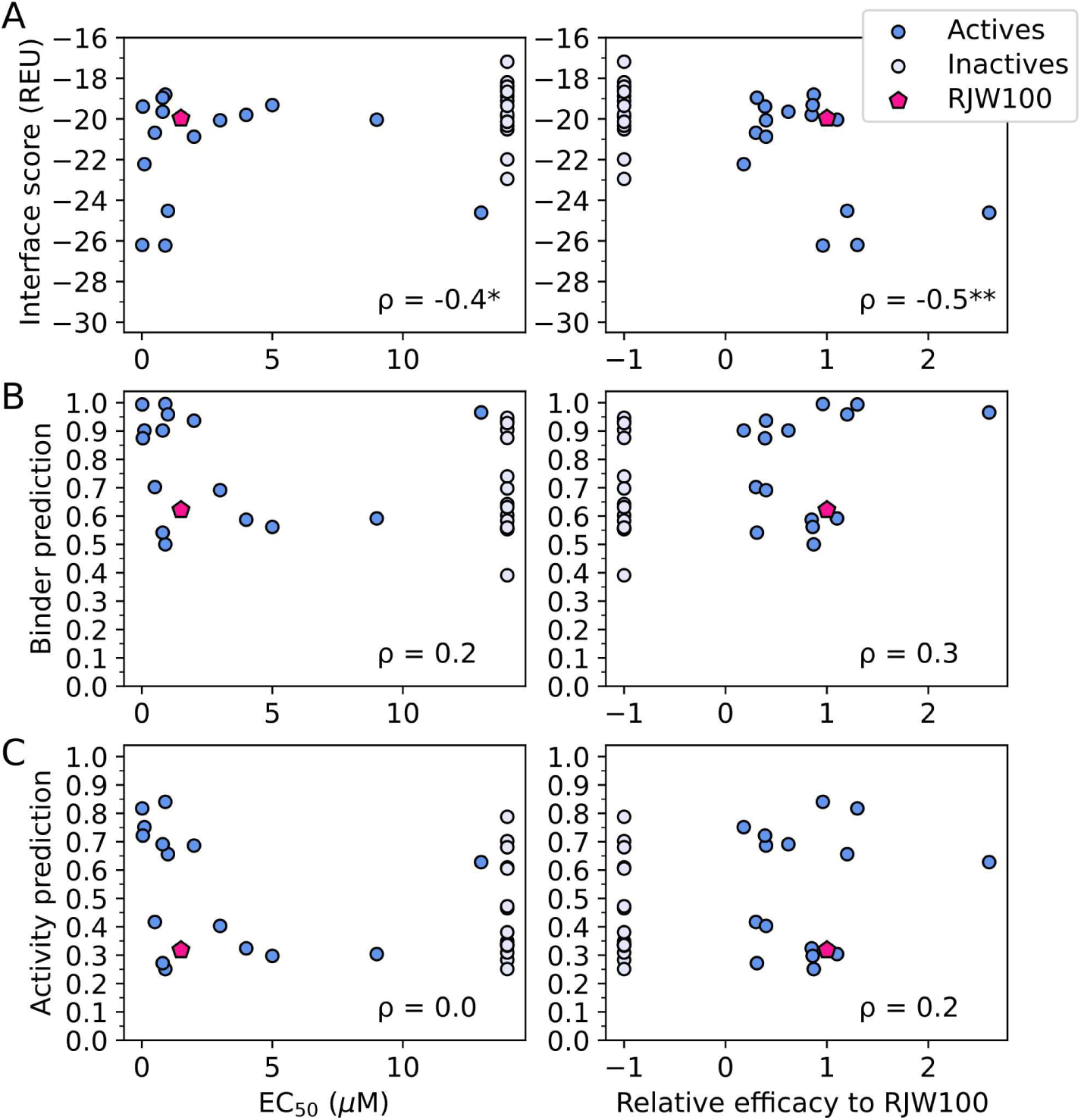
Correlation of computational metrics versus tested readouts for a congeneric compound series. (**A**) Interface score, (**B**) binder prediction from BLiP-L, and (**C**) activity prediction from ALiP-L versus EC_50_ (left column) or relative efficacy to RJW100 (right column) of the Mays and Mays_ia compounds. Values for EC_50_, relative efficacy, and activate/inactive labels were taken from reported data in Figure 2 of ^27^. Lower interface score indicates a more energetically favorable binder. Higher binder and activity predictions indicate that the models predict the compound is more likely to bind or activate LRH-1. Spearman’s rank correlation coefficient (ρ) calculated from the plotted data is reported at the bottom right of each panel, with statistical significance indicated by asterisks (*p-value < 0.05, **p-value < 0.01). No asterisk indicates a non-significant correlation.

**Figure S4.**
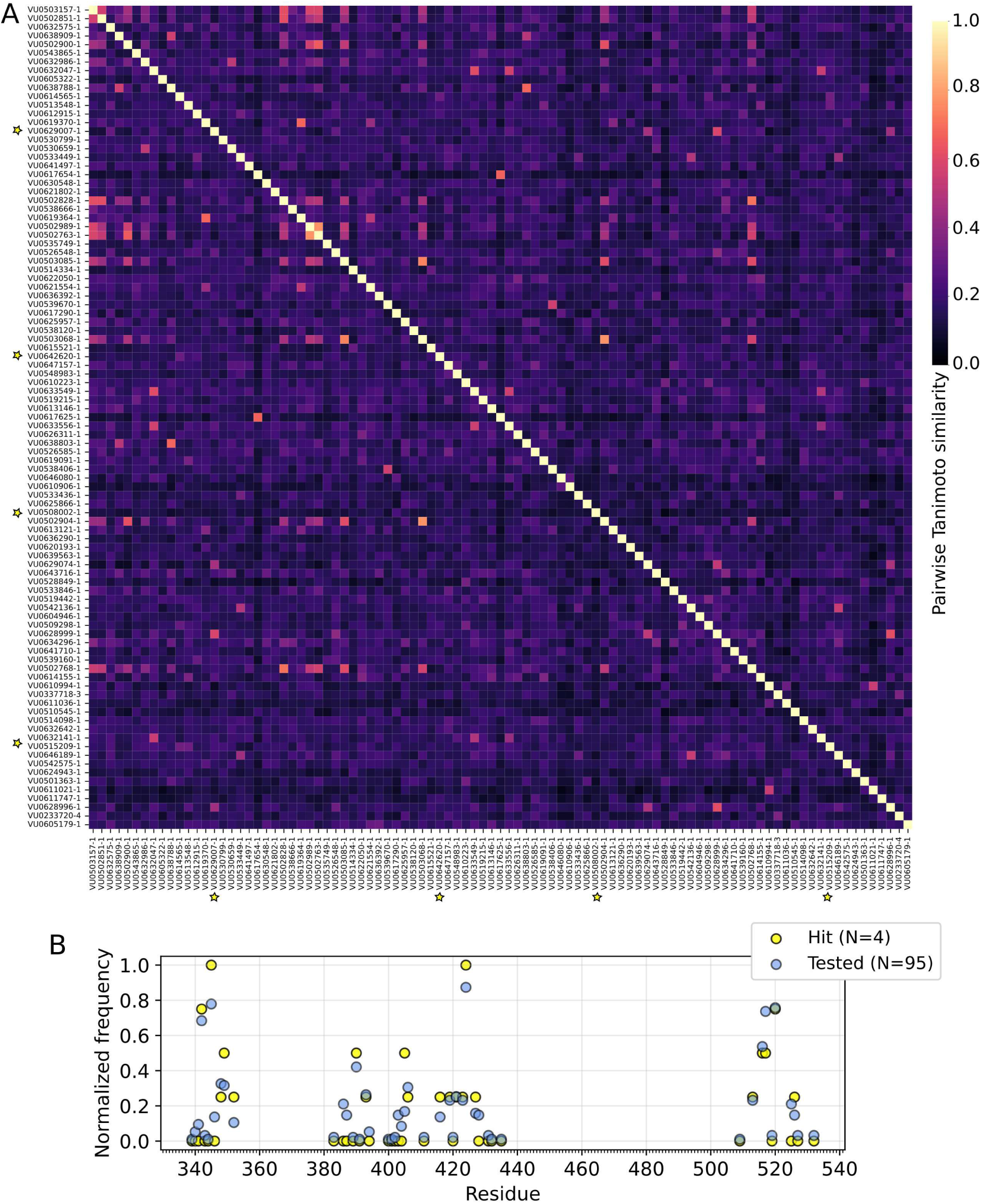
Profile of 95 selected compounds from VU98k library for testing. (**A**) Pairwise Tanimoto similarity between compounds computed using Morgan fingerprint. Values closer to 1.0 indicate higher similarity between compounds. VU identifiers reported for reference. Hit compounds from competitive binding assay are marked with a star. (**B**) Normalized frequency of ligand-LRH-1 residue contacts from lowest energy docked poses. Contacts were identified using CPPTRAJ native contact analysis and defined as non-hydrogen atom interactions within 3.0 Å between the compound and selected LRH-1 residues. Poses without contacts were excluded. Number of structures per label indicated in the legend.

**Figure S5.**
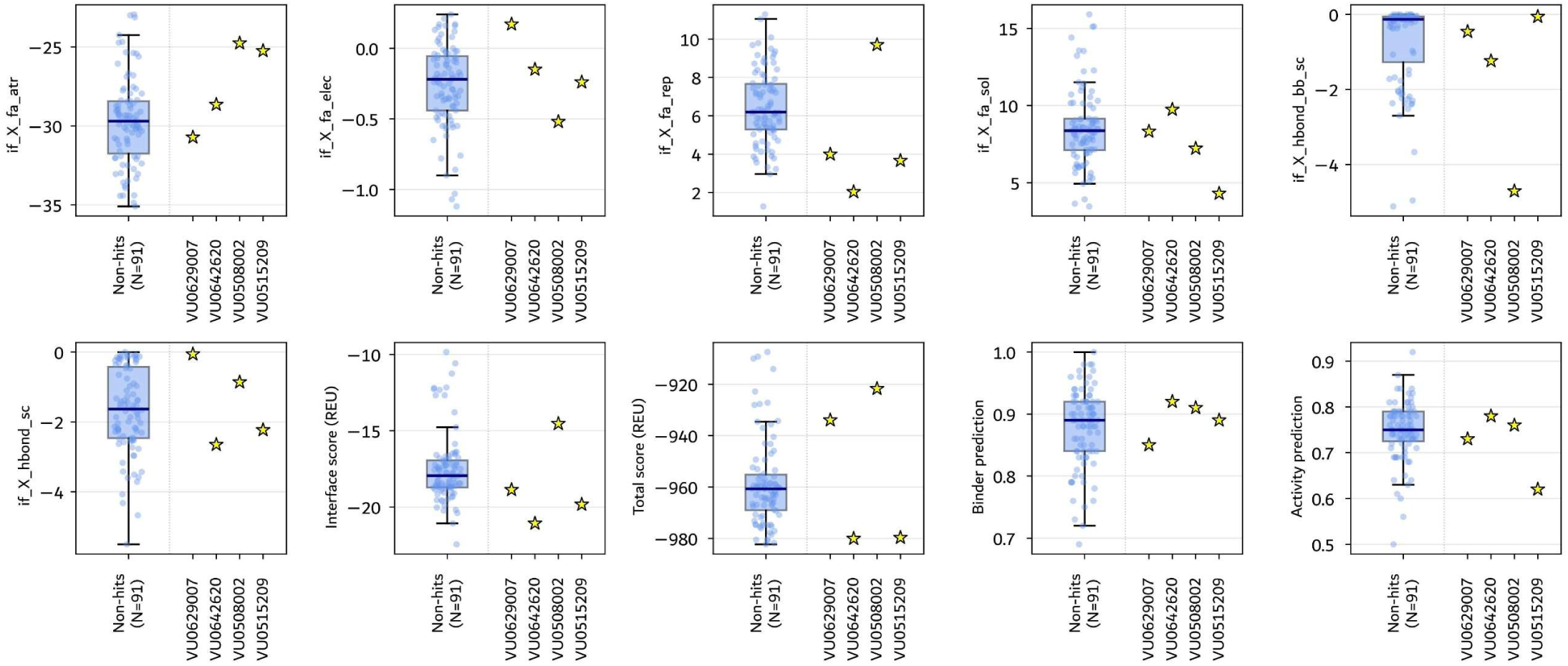
**Distribution of feature, score, and prediction metrics for tested compounds**. Box plots represent the distributions of the tested 91 non-hit compounds for the feature, score, or prediction metric specified on the y-axis label. Median is marked by the black line on the box plot, box bounds indicate interquartile range (IQR), and whiskers mark 1.5*IQR. Individual values are marked as blue scatter points. Yellow stars are individual values for the metrics of the hit compounds.

**Figure S6.**
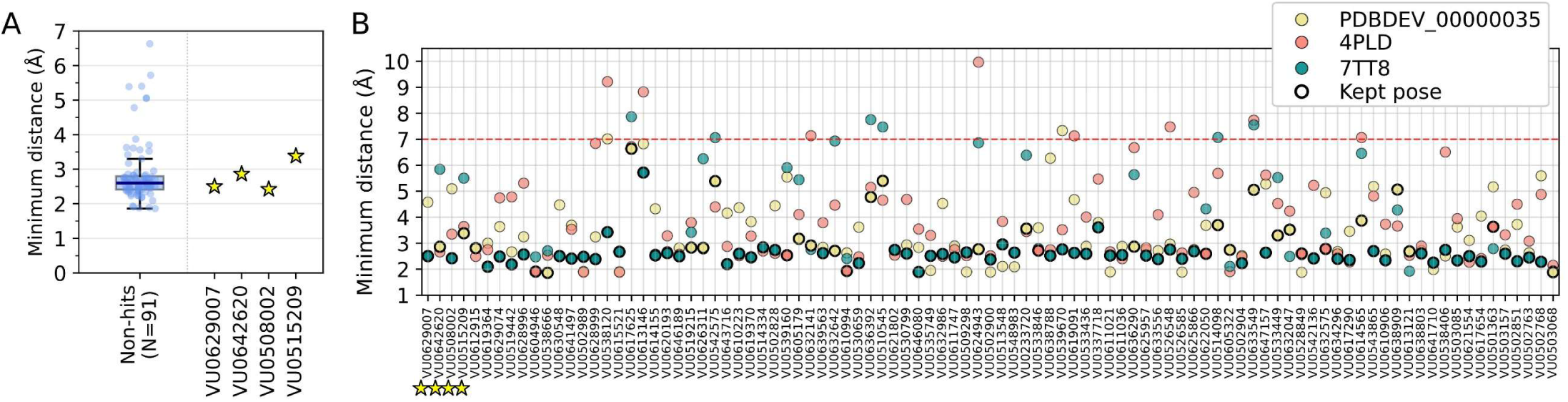
Minimum distance to predefined pocket residues for tested compounds. Minimum distance refers to the minimum heavy atom distance between LRH-1 residues G421, Y516, and K520 to any compound heavy atom in the docked pose. (**A**) Distribution of tested hit versus non-hit compounds. Non-hits box plot median is marked by the black line, box bounds indicate IQR, whiskers mark 1.5*IQR, and individual values are marked as blue scatter points. Yellow stars are individual values for hit compounds. (**B**) Minimum distance for the 95 tested compounds calculated from the lowest energy docked pose to each of the three pocket conformations. Hit compounds are marked by yellow stars. Colors correspond to the respective pocket conformation. Structure filtering cutoff threshold 7.0 Å marked by red dashed line. The pose considered after initial score and prediction metric filtering is marked by a thick black outline, note this pose may not always correspond to the lowest minimum distance pose.

**Table S1.** Percentage of tested hits and non-hits with checked chemical motifs.

| MOTIF | HITS (N=4, %) | TESTED NON-HITS (N=91, %) |
| --- | --- | --- |
| Carboxyl | 25% | 2.2 % |
| Ester | 25% | 7.7 % |
| Amide | 75% | 84 % |
| Ether | 0% | 11 % |
| Long chain C8 | 0% | 1.1 % |
| Chlorophenyl | 25% | 4.4 % |

**Table S2.** Key resources and reagents.

| REAGENT or RESOURCE | SOURCE | IDENTIFIER |
| --- | --- | --- |
| Bacterial and virus strains |  |  |
| BL21(DE3) *E. coli* | New England Biolabs | Cat# C2527H |
| Chemicals, peptides, and recombinant proteins |  |  |
| Cobalt Resin (HisPur) | Thermo Fisher | Cat# 89965 |
| Rh-PE Probe | Avanti Research | Cat# 810150 |
| 5Fam Coregulatory Peptides | Biosynth Peptide | Custom Order |
| RJW100 (Positive Control) | MedChemExpress | Cat# HY-131445 |
| Critical commercial assays |  |  |
| Capto Q Column | Cytiva | Cat# 17531601 |
| Deposited data |  |  |
| Human LRH-1 (NR5A2) LBD | UniProt | Prot: O00482 |
| TEV |  |  |
| Software and algorithms |  |  |
| Prism 10 | GraphPad Software | RRID: SCR_002798 |
| Other |  |  |
| 384-well black plates | Corning | Cat# 3573 |

## Materials and Methods

### Compound libraries

Method development and neural network model training used the 2322-compound Discovery Spectrum library (S2k), which is composed of 50% drug components, 30% natural products, and 20% other bioactive components. The library was previously screened against LRH-1 by Fluorescence Resonance Energy Transfer (FRET)^28^, where 58 unique compounds were identified to bind the orthosteric ligand-binding pocket in the LRH-1 ligand-binding domain (LBD). These 58 compounds were labeled as “binders” for model training, the remaining were labeled as “non-binders.” Four secondary assays identified 15 out of the 58 binders regulate LRH-1 function either by modulating coregulator peptide recruitment, the canonical mechanism of LRH-1 regulation, or impacting LRH-1 target promoters in mammalian cells^28^. These 15 compounds were labeled as “activators” for model training.

Prospective screening used the 98,765-compound Vanderbilt Discovery Collection (VU98k), a primary screening library housed by the Vanderbilt High Throughput Screening Facility composed of diverse drug-like compounds with lead-like motifs selected from Life Chemicals. Labels (binder, non-binder, activator, non-activator) are not known a priori to the screen.

Each library was processed as one structure-data file (SDF) containing all compounds in the library. The BioChemical Library (BCL)^54^ was used to assign 3D coordinates and generate conformers. Command-line usage of the BCL procedures were executed using custom Shell scripts for compound processing in a high-throughput format. For example, the following command results in 100 conformers for each of the input compounds:

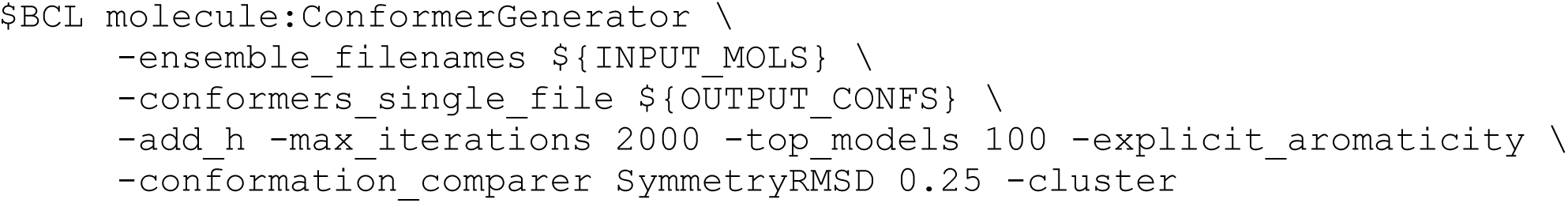

The command directs the BCL to perform 2,000 conformer-generating iterations by stochastic sampling of rotamer combinations from the crystallography open database while iteratively resolving clashes, score the conformers using a Boltzmann-like statistical energy function, then cluster and save 100 conformers based on score and filtered by structural similarity^74,75^. The saved conformers represent an energetically favorable and structurally diverse set to sample conformational flexibility of the compounds without input from the target pocket.

### Targeted docking to LRH-1 LBD pocket

Three LRH-1 models were selected to represent the LBD in three states: apo (4PLD^76^), holo with a synthetic small molecule bound in the target pocket (7TT8^12^), and as part of a full-length LRH-1 complex with a P6L phospholipid ligand bound in the target pocket (PDBDEV_00000035^77^). These models are nearly identical based on global conformation with minor local structural shifts in side chains for the residues near the pocket. Crystal waters and extraneous components (bound ligand, coregulator peptide, DBD), were removed leaving only the LRH-1 LBD. Missing residues were modeled using the Rosetta Remodel Mover^78^ such that all three constructs are the same length (240 residues). Renumbering ensured matching residue numbering, PyMOL (Schrödinger, LLC) was used to align the structures to 7TT8, and an all-atom Rosetta Relax with added constraints to backbone heavy atoms was used to refine the structures using the Rosetta force field^79^. Global RMSD indicates that the prepared LRH-1 LBD models were nearly identical (∼1 Å) and visual inspection confirmed the pocket remained open.

The decanoic acid tail of compound 6N-10CA (CCD IUW) from PDB 7TT8^12^ was removed to generate a modified reference ligand. The ligands from the processed screening libraries were aligned to the modified reference ligand based on the maximum common substructure using the BCL^54^ to place the ligands in the target pocket prior to docking. Rosetta params files storing the geometric and chemical information of each ligand were generated. The compound conformers and aligned compound were reformatted as PDB files with atom naming to match the params fille.

RosettaLigand with Rosetta XML scripting was used for docking^49–51,80,81^. RosettaLigand samples ligand conformers at the target pocket, defined by the aligned compound, allowing translation, rotation, and torsional adjustments of the compound and flexibility of the protein pocket with side chain repacking. Custom shell scripts were developed to execute RosettaLigand as follows:

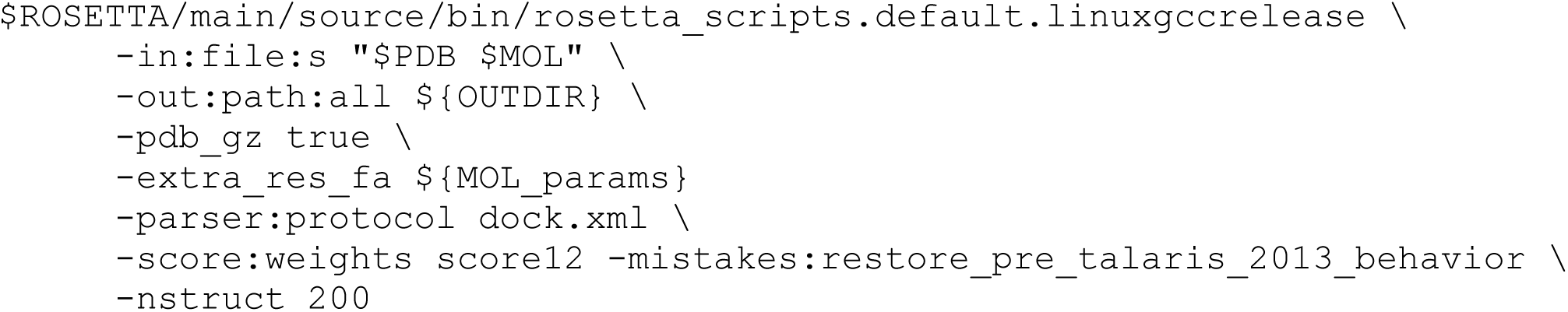

The prepared LRH-1 LBD models (variable named `$PDB`) and compounds from the screening libraries (variable named `$MOL`, param files and conformers defined by `$MOL_params`) were used as input. The docking protocol is defined in `dock.xml`, which is included in the accompanying GitHub repository for this manuscript. Standard docking protocols were referenced^50,51,80^. The Rosetta Transform mover was used to perform a Monte Carlo search of the binding site allowing up to 7.0 Å translation from the starting point of the compound in 0.2 Å increments with maximum 20° rotation angle per step followed by the Rosetta HighResDocker mover to perform side chain rotamer repacking. A cubical ligand scoring grid of 30 Å width was used to account for larger compounds. Use of the `score12` score function with `-mistakes:restore_pre_talaris_2013_behavior` flag reflects findings from a benchmark study demonstrating that this combination improves small molecule scoring^82^. RosettaLigand generates `-nstruct` docked poses per protein-ligand combination, each scored by the RosettaLigand scoring function and with individual energy terms reported.

### Blind docking against LRH-1 LBD

Blind docking was used to search for potential binding sites beyond the orthosteric pocket and explore alternative binding modes distinct from that of the aligned pose used in the targeted docking protocol. DynamicBind, a diffusion-based generative deep learning methods, was used for high-throughput blind docking for its command-line implementation and flexible treatment of the protein pocket^29^. DynamicBind was trained and validated using PDBbind2020^83^ with structures deposited before 2020, and therefore likely includes several LRH-1 protein-ligand complexes in its training set and would be biased towards predicting the orthosteric site as the binding pocket.

The same LRH-1 LBD models as prepared for targeted docking and the SMILES strings of the compounds were used as input to DynamicBind. DynamicBind outputs ten predicted poses per protein-ligand combination, ranked by predicted affinity score. The predicted poses were then scored using the RosettaLigand scoring function as done for the targeted docking models.

### Compiling select Rosetta energy terms for input features to neural network model

The Rosetta energy function includes terms to describe non-bonded interactions and physical and statistical potentials to account for bonds and torsional energetics which are combined to approximate the energy of a system^52,84^. The total score of a protein-ligand complex (total_score) is determined by a weighted linear combination of the individual energy terms and is reported in Rosetta Energy Units (REU). The interface score of a docked pose (interface_delta_X; IDX) is calculated as the difference between the bound protein-ligand state and a separated state where the ligand is moved away from the protein to remove all interactions^50^. A negative interface score therefore indicates favorable protein-ligand binding. The docked pose with the best (most negative) score is expected to be the most energetically favorable docked pose, likely capturing the native state, and therefore is typically used for ranking compounds in virtual screens.

The default weights in Rosetta score functions are optimized to correlate with experimental or structural data^52,85^. For example, the weights for the energy terms introduced in RosettaLigand were determined by maximizing the correlation between the weighed sum from generated docked poses and the native pose using 100 protein-ligand complexes from the PDB^49^. Six select Rosetta protein-ligand energy terms describe the van der Waals forces, electrostatics, desolvation penalty, and hydrogen bonding between a bound compound and protein (**Table 1**). These are denoted with “if_X” in the score file reported for each docked pose and extracted into a separate file to use as input features for binder and activity predictions. The reasoning for this is that select energy terms collectively describe the energetic profile of the LRH-1 protein-ligand binding interface and therefore can be used to train a model to predict binding. Translating this relationship to activity is less straightforward, as changes to activity involve allosteric or large conformational changes away from the binding interface. Protein-only energy terms were not included in the features because they will largely be similar between the LRH-1 complexes. In addition, differences due to changes in rotamer conformation for residues far from the binding interface could introduce noise to the model due to suboptimal configurations.

### Binder and activity likelihood model development

The architecture of prediction models BLiP-L (Binder Likelihood Prediction for LRH-1) and ALiP-L (Activity Likelihood Prediction for LRH-1) is a fully connected feed forward multi-layer perceptron (MLP) developed using the PyTorch v2.5.1 library. Both are binary classifiers trained using labels based on experimental data from the S2k compound library^28^. BLiP-L input features are the six Rosetta energy terms (**Table 1**) with a neuron-layer architecture of 6-24-6-1 (**Fig. 1B**). ALiP-L input features are the six Rosetta energy terms (**Table 1**) plus BLiP-L prediction logits with a neuron-layer architecture of 7-24-7-1 (**Fig. 1C**). For both models, the Leaky ReLU activation function was applied between hidden layers to support stable training, and output raw logits in the final layer. Training data included the Rosetta energy terms from protein-ligand docked models of the following. Dataset 1: all S2k compounds docked to target pocket in PDB 7TT8^12^ with minimum interface score less than zero. Datasets 2-4: the 58 binder compounds identified from a FRET-based screen on the S2k library^28^ docked to target pocket in PDB 4PLD^76^, 7TT8^12^, and PDBDEV_00000035^77^. Datasets 5-7: the 58 binder compounds blind docked against PDB 4PLD^76^, 7TT8^12^, and PDBDEV_00000035^77^.

This data augmentation technique samples the structural space of a compound with different LRH-1 LBD constructs and docking approaches. A total of two hundred decoys were generated for the full S2k docking (dataset 1), 4000 total decoys were generated for the 58 binder docking to each construct (datasets 2-4), and ten predictions were generated for the blind docking to each construct (datasets 5-7). The decoys were generated following the respective targeted and blind docking protocols detailed above. The decoy with the lowest interface score per dataset was used for training the models. This way, a binder compound may appear up to seven times in the training data, but the feature descriptors of the compound will be different in each entry. A non-binder compound only has one entry to avoid perpetuating data imbalance. Since even non-binders result in energetically favorable interface scores in Rosetta, a typical limitation of score functions, the sampling approach used in this work is intended to increase the chances of recovering a near-native pose with more energetically favorable decoys for the known binders. Without knowledge of the native pose of a compound, the sampled decoy poses provide a limited profile of plausible binding energy interactions for a given compound in the LRH-1 pocket. The total training dataset contained 2101 compounds, 359 entries labeled as binders and the rest labeled as non-binders.

Model development and hyperparameter tuning were conducted using a stratified 3-fold cross-validation scheme to maintain class distribution across splits with set random state reproducibility. Energy terms vary in scale, and therefore input features were standardized using Z-score normalization. The global mean and standard deviation were calculated from the training data only to prevent data leakage. A focal loss function was implemented to down-weight the contribution from easily-classified non-binders using a modulating factor *(1 − p_t_)^γ^*, where *p_t_* is the probability of a true class label and *γ=1* is a hyperparameter that controls how much the loss function down-weights clear false labels, to scale the standard binary cross-entropy loss. In this case, *γ=0* is standard binary cross-entropy loss. This prevents the majority non-binder labels from dominating gradient updates during backpropagation. A class-balancing weight ratio *N_neg_N_pos_* was used to ensure proportional contribution of the binder class. The model was trained using an Adam optimizer with learning rate 1e-4 for a maximum 3000 epochs. Early stopping was employed, which ends training if the validation loss fails to improve for 20 consecutive epochs to prevent overfitting and allow the model to generalize to unseen compounds. Following hyperparameter tuning, BLiP-L was trained using the total training set. The model architecture, weights, and normalization parameters, were saved for future use.

For ALiP-L, it was reasoned that any activator compounds should also be a binder, and therefore a feature informing on binder likelihood should be included. BLiP-L was used to generate binder likelihood values for the S2k compounds on which it was trained to include the binder likelihood feature for ALiP-L. While these in-sample predictions were subject to training set bias, the predictions were used solely as an additional input feature and not to assess model performance. To reduce data imbalance given the fewer number of activator compounds, the training data for ALiP-L excludes compounds with binder likelihood output logit below 0.23, which is the cutoff for the 70^th^ percentile of binder predictions. This way ALiP-L learns to differentiate between compounds that are predicted binders only compared to compounds that are activators. The training data for ALiP-L is reduced to a total of 630 compounds, where 67 compounds are labeled as activators. The same splitting, feature normalization, loss function, optimization, and training scheme were used for ALiP-L as detailed for BLiP-L. Learning rate was reduced to 1e-5. All other hyperparameters remained the same. The class-balancing weighed ratio for focal loss was automatically adjusted. The model architecture, weights, and normalized parameters were saved for future use.

Model performance was evaluated using both threshold-independent and threshold-dependent metrics. Area under the precision-recall curve (AUPRC) and area under the receiver operating curve (AUROC) were computed from prediction logits using the corresponding precision-recall and ROC curves. For binary threshold-dependent classification metrics, including accuracy and Matthews correlation coefficient (MCC), a fixed threshold was used where logits greater than zero were assigned to the positive class and therefore considered predictions for the positive class.

A permutation test where the features of the values in the training set were randomly scrambled was performed to evaluate whether the model predictions arose from meaningful feature-label patterns or random associations. Performance suffered for both binder and activity prediction models trained on jumbled features, with metrics indicating random guessing (**Fig. 1D,E**, *Trained model* versus *Jumbled*). This suggests that although the cross-validation metrics of BLiP-L and ALiP-L are not optimal, both models capture relationships between the energy terms and binding or activity labels. While there are no similar published models against which to directly compare BLiP-L and ALiP-L, the next section benchmarks BLiP-L and ALiP-L predictions against compounds with verified binding and activity against LRH-1.

### Testing trained models using Xtal and Mays compound datasets

An out-of-distribution test where the trained BLiP-L and ALiP-L models were used to predict binding and activity likelihood of compounds dissimilar to the training set S2k compounds was performed. Two datasets (“Xtal” and “Mays/Mays_ia”) were prepared for this test. To measure chemical similarity, compound SMILES strings were used to generate a chirality-aware 2048-bit Morgan fingerprint with radius 2, then the Tanimoto coefficient of the RJW100 Morgan fingerprint against each specified dataset compound was calculated using RDKit v2023.03.3 (**Fig. 2A**).

The “Xtal” dataset includes all compounds with published structural models, available at the time of the test, which bind to LRH-1 at the target pocket (**Table 2**). Three structures of the prototypical RJW100 compound were part of this dataset. To generate Rosetta energy terms from the crystallographic pose, crystal waters and extraneous components were removed and explicit hydrogens were added to the compounds using Open Babel^86^ v3.1.0, then Rosetta params files were generated. An all-atom Rosetta Relax with constraints to backbone heavy atoms was performed using the prepared protein file and compound as oriented in the structural model. Ten protein-only models were generated, the model with the best (most negative) total score was used for model prediction. The same RosettaLigand score function and weights as used to score the S2k compounds were used to determine the Rosetta interface score for Xtal compounds (**Fig. 2C**, *blue*) and corresponding energy terms were used for BLiP-L and ALiP-L (**Fig. 2D** and **2E**, *blue*).

The “Mays” dataset includes all compounds in Figure 2 of Mays et al^27^, which were synthesized and tested for binding to LRH-1 by measuring EC_50_ values and activity reported as relative efficacy to RJW100. Based on the nearly identical orientation of RJW100 to the crystallographic poses of 6N, 5N, and 2N analogs (**Table 2**), structural models for the Mays compounds were generated by performing in silico R-group substitutions on the RR-RJW100 from PDB 5L11. The BCL^54^ fragment mutate application was developed to maintain a starting compound binding pose while perturbing chemical groups in a drug design application. A detailed tutorial and example application of the protocol is in https://meilerlab.org/wp-content/uploads/2022/02/Tutorial_5.pdf. Its implementation into Rosetta as a mover eases use of this application into a pipeline to retain the input RJW100 binding pose while performing R-group substitutions.

A series of Shell scripts were developed to perform the following steps. First, SDF files of each individual substitution fragment were generated using RDKit. The attachment point for the fragment to the parent molecule was marked by an asterisk “*” to represent an unspecified atom. This is annotated as atom name “X” in the SDF. Next, the BCL `molecule:Mutate` application was used to remove the respective atoms for each of the three R-groups (R1 hydroxyl, R2 styrene, and R3 phenyl) while leaving the rest of the compound intact to generate an R-removed scaffold. Hydrogens were added to satisfy open valences. Rosetta params files were generated for each R-group parent. Then Rosetta with integrated BCL was used to append each modification to the respective R-group. The mutable atoms (R-removed scaffold) and fragment combinations were input to an XML script protocol which includes minimizing the scaffold in the target receptor pocket, performing design on the scaffold by appending the fragment to the scaffold using the `BCLFragmentMutateMover`, relaxing the designed molecule in the receptor pocket, filtering out poses with Rosetta total score above 0, and determining the interface score of the compound in the pocket. To generate both endo (N) and exo (X) diastereomers for the R1 substitution, the hydrogen bond removed to append a fragment was explicitly set using the `ov_shuffle_h` option in the `BCLFragmentMutateMover`. Five iterations of design (append fragment then relax modified compound in protein pocket) per R-fragment combination were performed. Scores and models were nearly identical given the intentionally limited sampling. A similar procedure as for the Xtal dataset was then applied to relax, score, and predict binding and activity for the Mays compounds (**Fig. 2C-E**, *red* and *white*).

### Prospective virtual screen of VU Discovery Collection library and compound selection for testing

Targeted docking was used to perform a structure-based virtual screen of the 98,765-compounds from the Vanderbilt Discovery Collection (VU98k) against LRH-1. Conformers were generated using the BCL as detailed in the “Compound libraries” section. As detailed in the “Targeted docking to LRH-1 LBD pocket” section, compounds were aligned to the modified reference ligand (6N-10CA from PDB 7TT8 without the decanoic acid tail) using the BCL followed by Rosetta params file generation. The same RosettaLigand docking protocol used to dock the S2k compounds was applied to dock the VU98k compounds to the target pocket of the LRH-1 LBD. Docking was performed against the three LRH-1 LBD constructs, one from an apo state and two from holo states with the compounds removed, using the same prepared PDB files as for the S2k compounds. To speed up docking, compounds were split into lists of 100 compounds and a SLURM job array was used to run RosettaLigand docking on the lists in parallel. For each compound-state combination, 200 docked poses were generated, the pose with the best (lowest) interface score was identified and its score and energy terms were saved for use in the prediction models. Compounds with only energetically unfavorable poses, indicated by docking scores above zero, were excluded.

BLiP-L and ALiP-L were used to get binding and activity likelihood predictions for the remaining compounds. Three sets of predictions per compound were computed based on the energy terms of the compound for each of the three LRH-1 LBD constructs. The best pose per compound as defined by the lowest IDX was used for the first round of filters, where compounds with sigmoid-transformed binder prediction equal to or less than 0.6 were excluded. Then the top 8000 compounds sorted by activity prediction and the top 4000 compounds sorted by IDX were saved. The cutoff and sorting steps were performed again for the compounds using the metrics computed based on the small-molecule bound holo LRH-1 LBD construct only. The filtered lists were combined, where for any duplicate compounds only one entry was kept.

Chemical properties were computed for the compounds in the reduced list, including molecular weight, number of heavy atoms, and LogP using Crippen’s method implementation in RDKit, and the generated compound SMILES string was saved. Compounds that did not pass at least two drug-like filters and which contained PAINS compounds were removed from the list to prioritize broadly drug-like candidates while tolerating minor deviations from a single criterion. The panel of drug-like filters checked were: Lipinski’s Rule of Five^87^, Ghose^88^, Veber^89^, Rule of Three^90^, REOS^91^, and a generic drug-like filter. Properties and cutoff criteria for the filters are detailed in the `utils_drugfilters.py` script. These scoring, predictive, and chemical filters resulted in a list of 17,106 entries from the 98k, out of which 1,962 compounds had two entries, one for the holo construct and one for a different construct.

The minimum atom-atom distance between any compound heavy atom and any heavy atom of three LRH-1 reference residues was calculated using the docked poses of the entries in the filtered list. The reference residues were G421, Y516, and K520. These pocket mouth residues are involved in direct hydrogen bond contacts with synthetic compounds 10CA, 6N, and 6N-10CA^12^, were found to maintain interactions with 6N-10CA in molecular dynamics simulations, and are implicated in allosteric communication pathways relating ligand binding in the pocket to the coregulator binding site^13^. Compounds with a minimum distance greater than 7.0 Å to all the reference residues were removed from the list. For duplicate compound entries, only the pose with the shortest distance to the reference residues was kept. This lenient filtering strategy was designed to incorporate guidance from published structures and structural studies into compound selection while accounting for uncertainty in docked poses. Structural filtering reduced the list to 11,290 (referred to the 11k list) unique compounds from the screened list of 98k.

Compounds were then assigned a confidence level based on predicted binding and activity probabilities. “High” confidence was assigned to compounds if both the sigmoid-transformed binder prediction was greater than 0.8 and the sigmoid-transformed activity prediction was greater than 0.7. “Medium” confidence was assigned if either prediction was greater than the high-confidence threshold and both predictions were greater than 0.5. Remaining compounds were assigned “low” confidence. Confidence thresholds were designed following guidance from benchmarking on the Xtal and Mays compound datasets (**Fig. 2**). This resulted in 5,423 high-confidence, 3,905 medium-confidence, and 1,961 low-confidence compounds.

Compounds were filtered to include at least one fatty acid mimetic fragment based on recent findings that these fragments are potential LRH-1 modulators^16^ and ordered by a top-50 ranking using ligand efficiency 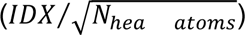 to normalize interface scores by compound size^92^. Compounds with identical ligand efficiency were assigned the same rank to avoid arbitrarily separating tied scores, resulting in more than 50 candidates. Predicted binding and activity probabilities were considered in parallel during prioritization but were not used as strict selection criteria. High-confidence compounds were prioritized for testing, while medium- and low-confidence compounds were also included to evaluate metric-based performance post experimental testing. The resulting subset was manually reviewed by visual inspection of docked poses, from which 80 compounds were selected for testing. An additional 15 compounds identified during earlier prioritization and manual review were retained despite later model updates that changed candidate rankings. Six of these 15 compounds did not contain fatty acid fragments. The final curated set of 95 compounds selected for testing included 68 high-, 23 medium-, and 4 low-confidence compounds.

### LRH-1 LBD protein expression and purification

The wild-type human LRH-1 (NR5A2) ligand-binding domain (LBD; residues 266–541, UniProt: O00482) and three Cys-lite mutants (C311/346S, C311/487S, and C346/487S) were cloned into a pET vector containing an N-terminal His-tag. Plasmids were transformed into *E. coli* BL21(DE3) cells and cultured in LB media at 37°C. Upon reaching an of 0.8, protein expression was induced with 1 mM IPTG, followed by overnight incubation at 15°C. Cells were harvested via centrifugation (6,000 x g for 30 min at 4°C) and resuspended in lysis buffer (20 mM HEPES pH 7.5, 150 mM NaCl, 2 mM CHAPS).Following lysis by probe sonication, the soluble fraction was isolated by centrifugation (12,000 x g). His-tagged LRH-1 was purified using immobilized metal affinity chromatography (IMAC) with Cobalt Resin. The His-tag was removed by TEV protease cleavage. Final purification was achieved via anion exchange chromatography using a Capto Q column, eluting with a linear NaCl gradient (0–1 M) in 20 mM HEPES (pH 7.5) and 2 mM CHAPS.

### Competitive binding screen with Rh-PE probe

Competitive binding was assessed using a Rhodamine-PE (Rh-PE) fluorescence polarization (FP) assay. Compounds were screened at 10 µM in 384-well black plates containing 100 nM purified LRH-1 LBD and 50 nM Rh-PE probe. RJW100 was utilized as a positive control, and wells containing only LRH-1 LBD and probe served as negative controls. Assay plates were incubated for 16 hours at room temperature under low-light conditions. FP was measured using a BioTek Neo plate reader (530/25 nm; 590/35 nm).

### Coregulator peptide binding assay

The recruitment of co-regulatory proteins was assed by polarization (FP) assay using 5-FAM-labeled peptides containing the LXXLL motif of DAX-1. Compounds were screened at 10 µM in 384-well black plates containing 0.38 µM and of purified LRH-1 LBD and 50 nM DAX-1 Fam labeled peptide. RJW100 served as a positive control, and wells containing only LRH-1 LBD and peptide served as negative controls. Assay plates were incubated for 16 hours at room temperature under low-light conditions. FP was measured using a BioTek Neo plate reader (Ex: 485/20 nm; Em: 528/20 nm).

### Intrinsic compound fluorescence interference screen

To identify potential false positives, the intrinsic fluorescence of the compounds was evaluated. Compounds were plated at 10 µM in 384-well black plates containing 50nM Rh-PE probe. Fluorescence intensity was measured at (Ex: 530.25 nm; Em: 590/35 nm) and compared to control wells containing only the Rh-PE probe. Compounds exhibiting significant intrinsic fluorescence relative to the probe-only control were flagged and noted during further high-throughput screen analysis.

### Statistical analysis

Statistical significance was determined using a one-way ANOVA with multiple comparisons to the control wells. Data were log_2_-transformed prior to analysis. All statistical tests were performed using GraphPad Prism software.

## (Supplementary Material)

## Acknowledgments

Experiments were performed in the Vanderbilt High-Throughput Screening (V-HTS) Core Facility with compound management assistance provided by Corbin Whitwell. Vanderbilt’s Advanced Computer Center for Research and Education (ACCRE) compute cluster was used for high-performance computing needs including virtual screening. Vanderbilt’s Center for Structural Biology manages the hardware used for model training and data analysis.

## Author Contributions

ACCG was the primary author of the manuscript, including developing the code for training, testing, and analyzing the machine learning model, running benchmarks, generating figures, and writing text. ANC performed all wet lab experiments and authored the related methods sections. EWB provided guidance on model development, experimental design, data analysis, and overall research direction. JM and RDB oversaw the research and provided intellectual guidance. All authors assisted in and approve this manuscript for publication.

## Data and Materials Availability

All data needed to evaluate the conclusions in the paper are present in the paper, the Supplementary Materials, and/or in https://github.com/meilerlab/lrh1-binding-activity-prediction.git. The files in the repository in their current form will be archived in Zenodo after publication.

